# Recent horizontal transfer of transposable elements in *Drosophila*

**DOI:** 10.1101/2025.10.30.685650

**Authors:** Shashank Pritam, Sarah Signor

## Abstract

Transposable elements are small pieces of DNA which selfishly increase in copy number within genomes. Transposable elements can also move between species through unknown intermediaries in a process known as horizontal transfer, infecting novel genomes and increasing in copy number. In this manuscript we used almost 400 dipteran genomes to uncover 648 recent transposon invasions, mostly in *Drosophila*. We limited our results to transposons with 99% similarity to capture recent transfers. When our results overlapped with previous studies they were largely replicated. The majority of transfers occurred between closely related species, with the cosmopolitan *melanogaster* group showing the highest recent transfer activity. However, we documented 60 HT events spanning more than 30 million years of evolution. *Gypsy* and *Mariner* elements were involved in the most HT events. While the majority of elements were implicated in fewer than five transfers, we documented a single transposon with as many as with 16 recent transfers, many between different *Drosophila* groups. In addition, we found that while LTR elements engage in HT more frequently, DNA elements travel longer phylogenetic distances when engaging in HT. This potentially represents a different evolutionary strategy for exploiting naive genomes. Our phylogenetic framework advances the understanding of horizontal transfer dynamics at the species level within *Drosophila*.

## Introduction

Transposable elements (TEs) are mobile DNA parasites which increase in copy number within their hosts’ genome. They frequently interrupt functional sequences, cause double stranded breaks, and induce ectopic recombination. Recently, it was demonstrated that in *D. melanogaster* as much as 20% of recessive lethals are transposon insertions [Marion et al., 2026]. Due to this and other negative side effects, host species have specialized systems for suppressing the movement of transposons [Brennecke et al., 2007, Aravin et al., 2007, Brennecke et al., 2008]. In *Drosophila*, this system relies on specialized small RNA from the PIWI-interacting RNA (piRNA) pathway [Senti and Brennecke, 2010, Brennecke et al., 2007, Gunawardane et al., 2007]. These piRNA are cognate to the TE, and can then be used to target TE transcripts for degradation, or TE sequences for silencing [Brennecke et al., 2008, Teixeira et al., 2017, Aravin et al., 2007]. piRNA are typically generated from TE copies inserted within specialized genomic regions, known as piRNA clusters [Brennecke et al., 2007, Malone et al., 2009]. This system adapts to the invasion of a new TE through an as-yet not fully understood mechanism. In the germline, the TE will either insert into an established piRNA cluster such as *42AB* in *Drosophila*, or will begin producing piRNA *de novo* [Srivastav et al., 2023]. While TE silencing is not absolute, there is evidence that over time selection is lessened on the TE and the TE sequence will begin to degrade. One way in which TEs can escape this fate is by transferring to the naive genomes of other species, or horizontal transfer (HT) (Figure 1) [Kofler et al., 2015a, Scarpa et al., 2025, Pianezza et al., 2023].

**Figure 1.**
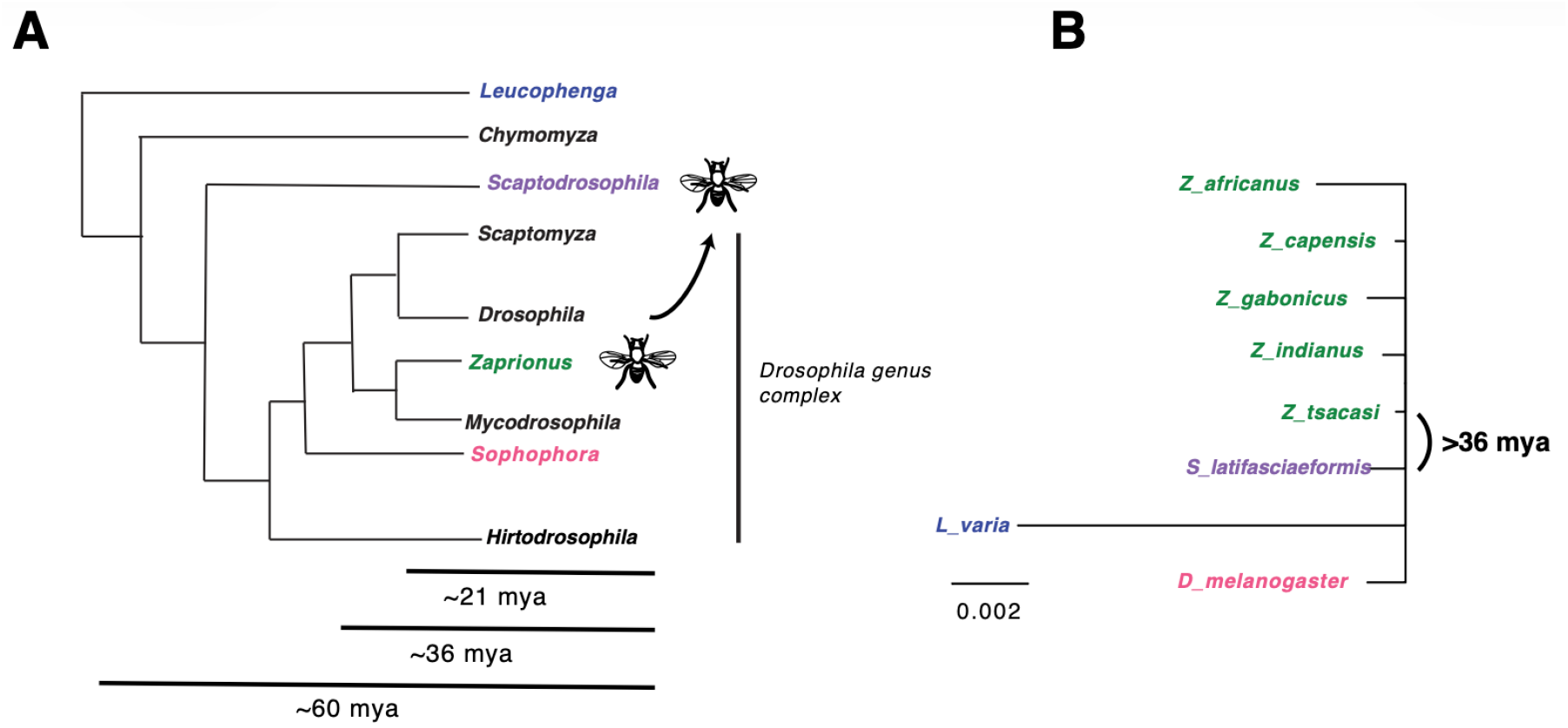
An illustration of horizontal transfer in the Drosophilidae. **A.** Phylogeny of the Drosophilidae, including approximate divergence dates of several nodes on the tree. The group as a whole likely began diversifying 60 mya, while the split between *Drosophila* and *Sophophora* is closer to 36 mya. Estimates are drawn from [Dias et al., 2025]. **B**. An example of a TE which has moved between distant members of the Drosophilidae. The tree is a star phylogeny with each sequence more than 99% similar, yet for example *Zaprionus* and *Scaptodrosophila* likely diverged more than 36 mya.

HT of TEs has been documented extensively for a limited number of TEs. For example, the transfer of the *P*-element from *D. willistoni* to *D. melanogaster* was discovered due to hybrid dysgenesis between *D. melanogaster* strains from different locations [Daniels et al., 1990]. Hybrid dysgenesis occurs when a TE is present in the male parent and not the female, and typically causes fertility defects. The *P*-element was also found to have transferred from *D. melanogaster* to *D. simulans* sometime after 2005 [Kofler et al., 2015a]. The discovery of these invasions is due in large part to the intense sampling to which these species have been subjected. More recently, sequencing of museum specimens has expanded our understanding of the invasion history of *D. melanogaster*, as has the availability of *D. melanogaster* datasets from populations around the world [Scarpa et al., 2023, Pianezza et al., 2023, Pianezza and Kofler, 2025]. This work identified up to 11 TEs which transferred to *D. melanogaster* in the last 200 years, a far greater rate than previously expected [Scarpa et al., 2023, Pianezza et al., 2023, Pianezza and Kofler, 2025]. However, this is a relatively small dataset with which understand broader patterns of HT.

Broader patterns of HT have been documented across insect lineages, with comparisons between Hemiptera, Hymenoptera, many lineages of diptera, and other groups such as Coleoptera [Peccoud et al., 2017]. This broad scale study found that *Mariner* elements were responsible for the most HT events, followed by *Helitrons* and *hAT* elements [Peccoud et al., 2017]. *Mariner* elements were noted as moving larger phylogenetic distances than retrotransposons, though the vast majority of insects in the study had diverged more than 100 mya. Furthermore, the transposons in the study are not necessarily highly similar so much as they are more similar than expected by the neutral mutation rate. This makes a good case for the conclusion that *Mariner* elements may be less dependent on host factors when transferring compared to other TEs. However, a comprehensive dataset of fine-scale HT is still lacking.

While the phenomenon of HT has been clearly documented, its mechanism is not. One of the more widespread hypotheses is that viruses facilitate the transfer of TEs [Miller and Miller, 1982, Fraser et al., 1985, Jehle et al., 1995], but parasitoid wasps and parasitic mites have also been implicated in HT [Yoshiyama et al., 2001, Houck et al., 1991]. *Wolbachia*, an intracellular parasite that is widespread in insects, has also been considered a possible vector [Kondo et al., 2002, Heath et al., 1999]. None of these vectors has been demonstrated to mediate the transfer of a TE in laboratory or natural settings. One class of TEs (*gypsy/gypsy*) contains an *envelope* protein which allows the TE to act like an endogenous retrovirus and gives them infectious properties (*ZAM, springer, Tirant, Quasimodo*) [Song et al., 1994, Senti et al., 2025]. However, in *Drosophila*, it is only one family of TEs that possess *envelope* proteins, and HT is observed for all classes of TEs regardless of the presence of this protein. Some TEs do appear to engage in HT more frequently than others - for example *Mariner* elements have been documented crossing large phylogenetic distances in *Drosophila* [Peccoud et al., 2017, Maruyama and Hartl, 1991a, Robertson, 1993]. However, depending on the study, it is not always clear if it is the same TE or a member of the same class of TE.

In order for a TE to spread between species it must either come into contact with a host or with a vector from the original host species. This generally requires ecological proximity [Venner et al., 2017]. TEs are also much more readily transferred between more closely related species [Feschotte and Pritham, 2007, Peccoud et al., 2017]. Thus, for example, many of the recently acquired TEs in *D. melanogaster* are transferred from *D. simulans*, a close relative that is also a cosmopolitan human commensal which occupies many of the same ecological niches as *D. melanogaster* [Kofler et al., 2015a, Scarpa et al., 2023, Pianezza et al., 2023, 2025, Scarpa et al., 2025]. However, ecological proximity is not correlated with phylogenetic proximity [Peccoud et al., 2017]. As humans move organisms and goods, the rate at which organisms experience novel congeners will also increase. A not yet appreciated byproduct of this contact between congeners could be the exchange of novel TEs [Scarpa et al., 2025]. The expectation is that the transfer of TEs may have accelerated along with human commerce and transport. What the implications of that may be for genomic stability are unknown. For example, in *D. melanogaster* recent TE invasions have increased its genome size by 1 MB [Pianezza et al., 2025].

In this manuscript, our aim is to investigate fine scale HT in *Drosophila*, limiting ourselves to recent invasions. We were interested to see the rate of horizontal transfer of TEs, the species between which transfer was occurring, and the phylogenetic distance between donor species. We found evidence of more than 600 horizontal transfers within *Drosophila* and other closely related dipterans such as *Zaprionus* and *Lordiphosa*. We recapitulated previous results, such as the invasion of the *P*-element and *Spoink*, as well as many novel transfers. Some of our results were expected, for example, the majority of HT events were found in LTRs, which are known to be the most active TEs in *Drosophila* [Kofler et al., 2015b, Mérel et al., 2020]. Other results were less expected, such as the observation that DNA transposons transfer less but cover larger phylogenetic distances than LTR transposons. When HT of insect TEs has been examined at large scales, *Mariner* elements were most frequently involved in HT. Retrotransposons and DNA transposons likely have separate evolutionary strategies - at short phylogenetic distances retrotransposons transfer most frequently, while at larger phylogenetic distances DNA elements are the most active.

## Methods

### Genetic resources

All *Drosophila* genomes used in this manuscript were obtained from NCBI and previously published [Kim et al., 2023, 2021]. The TE library was obtained from [Chakraborty et al., 2021]. This TE library contains 2, 332 repetitive elements from a variety of species groups, including *grimshawii, ananassae, willistoni*, and *ficusphila*. We excluded satellite repeats and *R* elements (which are vertically inherited)[Eickbush and Eickbush, 1995] as well as redundant TEs.

### Identifying horizontal transfer

TE insertions were initially located using the entire library of TEs and genomes with blastn [Camacho et al., 2009]. These files were filtered for alignments of at least 80 percent similarity to the reference and 1000 bp in length. To obtain recently active TEs, they were then filtered for hits that were at least 80 percent of the reference length for the TE internal sequence. To obtain a consensus sequence of each TE from each species, the sequence of each blast hit was extracted and aligned with mafft [Katoh et al., 2002, Katoh and Toh, 2008, Katoh et al., 2009]. For each species, a consensus sequence was created with EMBOSS consensus [Rice et al., 2000]. Following this, the sequences from each species were aligned with mafft [Katoh et al., 2002, Katoh and Toh, 2008, Katoh et al., 2009]. To verify the results of the consensus sequence approach, we also created a dataset of aligned individual insertions using the same basic pipeline but without the creation of a consensus sequence (see Supplemental File 1 for examples).

Alignments were hand-checked for errors. TEs present in three species or less were excluded from the dataset. Alignments were used as input to MrBayes [Ronquist et al., 2012, Huelsenbeck and Ronquist, 2001]. The phylogenetic trees were built using a GTR substitution model and gamma distributed rate variation across sites. The Markov chain Monte Carlo chains were run until the standard deviation of split frequencies was below 0.01. The consensus trees were generated using sumt conformat = simple. While constructing branch lengths, MrBayes uses evolutionary models for nucleotide and branch lengths that are supposed to represent the expected number of substitutions per site [Nascimento et al., 2017]. A tree for each TE was constructed in R using the package ape [Paradis et al., 2004, Paradis and Schliep, 2019].

### TE and species similarity

CLUSTAL was used to generate distance matrices from each of the aligned TE sequences [Sievers and Higgins, 2018]. We selected TE alignments that were *>* 99% similar in different species as recent invaders. This is an arbitrary threshold, however, it does guarantee that any TE in the dataset was likely recently active. To be sure this represents an HT event and not interbreeding or recent speciation, we excluded species that were not at least 98% diverged. We used sequences from *Adh, Gpdh*, and *Ddc* to gather the percent divergence for species pairs in the dataset, see (Supplemental File 2). This excluded transfers between species such as *D. saltans* and *D. austrosaltans*, or *D. rhopaloa* and *D. carrolli*. At times these species are treated separately - for example if a TE is present only in *D. carrolli* at *>* 99% but not in *D. rhopaloa*. Otherwise, they were treated as a single unit. We also excluded all Hawaiian *Drosophila* due to their close relatedness and recent origin. Note that if a TE is active in two different groups, i.e. *Zaprionus* and *melanogaster*, it is not necessarily *>* 99% similar between the two groups. However, to ensure that we are annotating the same TE, we followed the 80-80-80 rule, such that they are 80% similar for 80% of their length (Supplemental File 3). Whether two TEs that are 80% similar and active in *Zaprionus* and *melanogaster* are the same TE is a philosophical matter for another manuscript. However, please see Supplemental File 3 for a discussion of the impact of this rule upon the dataset (which is quite minor).

### Manual curation of the HT dataset

Due to the difficulty in making phylogenetic inferences about HT, we manually curated a data set using the HT events described above as our starting point. It is quite difficult to infer the direction of transfer of a TE when it is *>* 99% similar. In terms of which TE transferred to who our dataset is direction agnostic. When transfer between more than two species that are all *>* 99% similar was noted, we used the branch length and phylogenetic position assigned by MrBayes to assign transfer partners. This may be somewhat arbitrary, but the majority of our analysis is not based on the exact donor species within a group. Note that if the route by which a TE transferred was unclear - i.e. equal similarity between *D. bipectinata, D. pseudoananassae, Z. inermis*, and *Z. nigranus*, we took the most parsimonious route. In this example, a transfer would occur between *D. bipectinata* and *D. pseudoananassae*, followed by a single transfer to either of these species to the *Zaprionus* species, and a transfer between the two *Zaprionus* species. Essentially, intra-group transfers were favored over inter-group transfers, and inter-group transfers were favored over inter-genera transfers (Supplemental File 4). Because of this, the majority of our analysis is at the level of subgroup or subgenus.

## Results

### TE Dataset

TEs less than 1000 bp were removed from the dataset, as were *R* elements and satellite repeats. The final dataset included 1410 TEs. After blasting to the *Drosophila* assemblies we removed TEs found in three species or less. We blasted each TE dataset to itself to remove any redundant hits and confirm that they were reciprocal best blast hits. We identified 648 invasions with *>* 99% similarity between species (Supplemental File 4). The majority of these invasions occurred within the *melanogaster, ananassae, rhopaloa*, and *willistoni* (sub)groups (fig. 2A). TEs are frequently active in different groups, and therefore at some time a transfer between groups may have occurred, but it is only included in the dataset if the TEs are *>* 99% similar between species. In general, we assumed the most parsimonious explanation for TE transfer if the exact transfer partner was not clear - we favored transfers within groups and within genera. For example, if a TE transferred from *Drosophila* to multiple species of *Zaprionus*, we assumed a single transfer from *Drosophila* followed by within-group transfers rather than multiple transfers from *Drosophila* (fig. 2). The only time this rule was not followed is if the more phylogenetically disparate pair was significantly more closely related than the less disparate pair.

**Figure 2.**
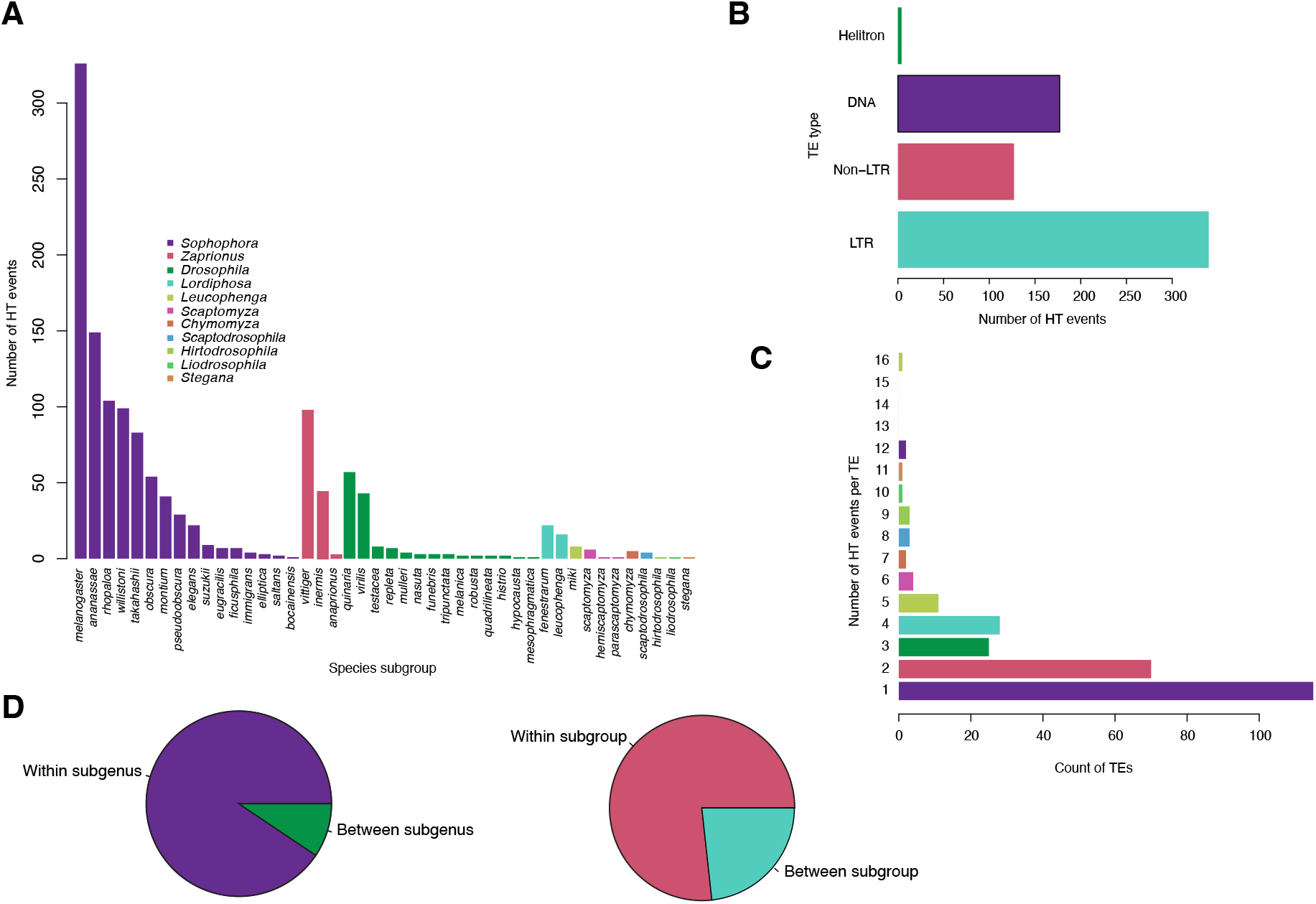
HT at the level of subgenus and subgroup. **A** The number of TE transfers for each subgroup in the dataset. This is agnostic to the direction of transfer and includes both partners. The distribution of this graph is largely determined by the number of species in each subgroup included in the dataset and the number of annotated transposons, rather than by any biological differences. **B** The number of TE transfers attributable to each type of TE in the dataset. **C** The number of TEs with each documented number of transfers. The majority of TEs are documented as only a single transfer, and at the other extreme a single TE recorded in 16 transfers (*Minona*). **D.** Summary of the number of HT evens that occurred within subgenera and subgroups. The number of species included outside of Sophophora is considerably less than within, so the dataset will be biased towards transfers within Sophophora.

The total number of TEs involved in HT is 268 (for 648 invasions), 47% of which are documented in a single transfer (Figure 2C). Of the 268 TEs included in the HT dataset, 57% of these TEs are LTRs, 22% are DNA elements, and 21% are Non-LTRs (Figure 2). Given that the dataset consists of TEs curated primarily from *Sophophora*, the majority of transfers were recorded between species of *Sophophora*. However, *Zaprionus* and *Drosophila* are tied for second most frequent (Figure 2A).

### Horizontal transfer of transposable elements

The most common TE types to be involved in an HT were *BEL, Mariner*, and *Gypsy* (Figure 3). *Mariner* has previously been recognized to be involved in frequent HT between species [Maruyama and Hartl, 1991b,a, Wallau et al., 2014, Robertson, 1993]. The vast majority of HT events occurred between closely related species within the same subgroup (Figure 2D). However, 151 of the transfers occur between different subgroups (i.e. the *melanogaster* or *ananassae* species subgroup). 61 transfers occurred between different subgenera, such as between *Zaprionus* and *Drosophila* (Figure 2D). Furthermore, 35 TEs are active in more than one group, but would not have been included in the count of subgroup transfers because the transfer between groups is not documented. Including different transfers under a single TE is heavily affected by how a TE is defined, which here is through 80-80-80 rule [Wicker et al., 2007]. In the entire dataset, if we raised the similarity requirement to 90%, 18 TEs would be split into two TEs. In addition, if additional species pairs are excluded based on potential for vertical transfer (i.e. *D. simulans* and *D. sechellia*), the number of transfers is reduced to 570 though the broader patterns are unchanged (Supplemental File 5).

**Figure 3.**
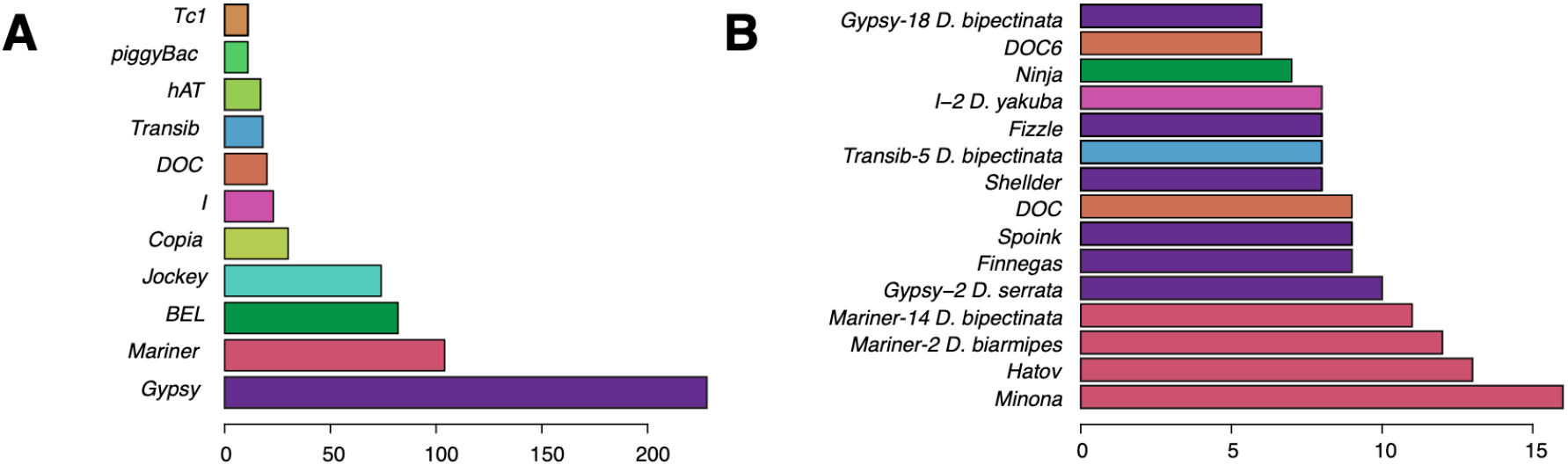
HT patterns among different TE groups. A. The frequency of HT observed for each type of TE. B. The 15 TEs with the most HT events in the dataset. Bars are colored according to the superfamily membership shown in the previous figure, i.e. dark pink indicates a Mariner element.

### Validation of the approach

*D. melanogaster* is the only species in which recent HT has been comprehensively documented. TEs that are known to be involved in HT based on museum specimens or direct observation include *412, I*-element, *opus, P*-element, *Spoink, Tirant, Hobo*, and *Blood. Blood, 412, Spoink, I*-element, *opus, Tirant*, and *Hobo* are more than 99% similar between *D. melanogaster* and *D. simulans*, confirming previous results from museum specimens documenting HT between the species (Figure 4B). The *P*-element is also confirmed in this dataset as moving from *D. willistoni* to *D. melanogaster* and from *D. melanogaster* to *D. simulans* (Figure 4D). The transfer of the *P*-element is one of the most well documented HT events in the literature.

**Figure 4.**
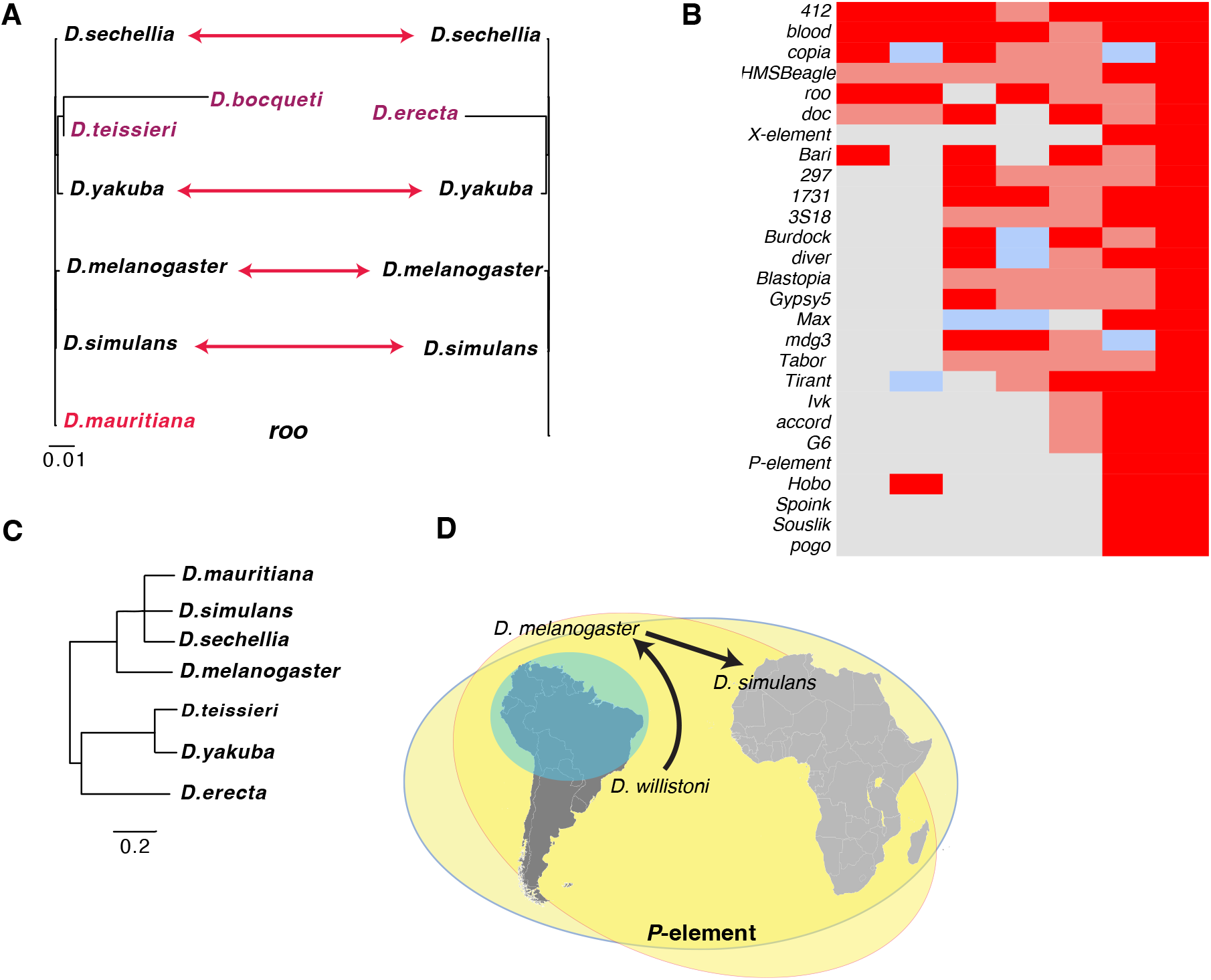
Validation of HT detection methods. A. The predicted phylogeny of *roo* in this study (left) compared to de la Chaux and Wagner 2009. Species in purple differ between the studies (i.e. de la Chaux and Wagner did not include genomes of *D. boqueti* or *D. teissieri. D. mauritiana* is shown in red because it was not included in the 2009 study but does have an additional HT event in this dataset. B. A chart of Tes that were investigated in: This study (2025), Pianezza (2025), Modolo (2014,) Carereto (2011), Bartolome (2009), Loreto (2008), and Sanchez (2005). Note that dark red indicates a perfect match, while light red indicates that one more or one less species was included. For example, if we concluded the transfer pattern was either *D. simulans* or *D. sechellia*, and another study listed only *D. simulans*, it would be marked light red. Gray indicates it was not included in the manuscript, and blue indicates a different species. Many of the differences are due to differences in the goal of the study - for example Pianezza (2025) wanted to identify the transfer partner of *D. melanogaster*. This could have happened in the deeper past, with a recent transfer to *D. simulans* that would be included in our dataset.) C. The species phylogeny of the *D. melanogaster* group, reconstructed from Seetharam (2013) [Seetharam and Stuart, 2013]. Note the difference in scale between A. and C. in terms of divergence, and the change in phylogenetic relationship of *D. yakuba* relative and *D. teissieri* in the *roo* phylogeny. D. The invasion path of the *P*-element as reconstructed in this manuscript and previous work. The blue circle indicates the range of *D. willistoni*, while the two yellow circles indicate the cosmopolitan range of *D. melanogaster* and *D. simulans. Gypsy-30 D. willistoni* is also among the most active TEs, though only two transfers occurred in the *willistoni* group. Rather, this TE is active in the *obscura, melanica, virilis, robusta, saltans, willistoni*, and *zaprionus* (sub)groups. While the transfer to *zaprionus* is not included in the dataset, it most likely came from the *melanogaster* group. Decreasing the threshold to 98.5% increases the number of transfers by five, to include two more members of the *willistoni* group (*D. paulistorum* and *D. equinoxialis*, as well as *D. tripunctata, D. putrida*, and *D. americana*. This would increase the number of subgroups in which the TE is active as well, to include *tripunctata* and *testacea*.This demonstrates the difficulty with naming TEs after a number and species, as they are typically active in groups outside of that species, if they are active in the named species at all. Copies of this TE are more than 90% similar between groups. Typically copies are more than 97% similar, with those between *Zaprionus* and *Drosophila* subgroups between 92%-94% similar. We propose to name this TE *Finnegas*.

Several manuscripts have also documented the presence and potential HT of *roo*, a particularly abundant TE in *D. melanogaster* (Figure 4A) [Modolo et al., 2014, de la Chaux and Wagner, 2009]. In Modolo (2014) and de la Chaux (2009) *roo* was inferred to have moved between *D. simulans, D. sechellia, D. yakuba*, and *D. melanogaster* (Figure 4A). Aside from methodological differences, our results are entirely in agreement with previous studies. For example, the original manuscript on *roo* did not include several species that we included, such as *D. teissieri, D. bocqueti*, and *D. mauritiana. D. mauritiana* is the only species with an additional HT event compared to the original manuscript. The species tree is shown in (Figure 4C), illustrating that the TEs between species are much more closely related than expected, and *D. yakuba* and *D. teissieri* do not follow the expected phylogenetic patterns.

Other TEs have been investigated in the literature, and our results are concordant with their findings. A literature review including Pianezza (2025), Carereto (2011), Loreto (2008), Modolo (2014), Bartolome (2009) and Sanchez (2005) of TEs in *D. melanogaster* found 27 HT events that are shared with our dataset (Figure 4B). Overall, 24/27 HT events were the same between essentially all of these datasets. There was significant disagreement for *Copia* and the *Max*-element. *Copia* originated from either *D. simulans, D. sechellia, D. yakuba*, or *D. ercepeae*, depending on the study, and our work is consistent with *D. simulans*. In Bartolome (2009)and Carereto (2011) the *Max*-element was thought to have transferred from *D. yakuba* to *D. melanogaster*, while our work is consistent with Pianezza (2005) which concluded it originated from *D. simulans*. Other research that confirms our results includes *POGO*, which was not previously found in any *melanogaster* group species except for *D. melanogaster*. This is recapitulated in our dataset where it is found in *Zaprionus, Leucophenga, Scaptodrosophila*, and *D. melanogaster*. There are undoubtedly significant false negatives in our dataset - our cutoffs are strict despite a TE with 98.5% divergence likely also being a very recent transfer. However, our goal was not to discover every HT, but to compile a large and accurate dataset of HT. Overall, these are relatively minor differences and the bulk of our analysis is consistent with previous research.

### TEs with the most horizontal transfer events

The top five TEs involved in HT include 4 *Mariner* elements and one *gypsy* element. 6 of the top 15 HT TEs are *gypsy*-class elements, while *Doc* and *I-2* are Non-LTRs. *Transib-5 D. bipectinata* is a DNA element like *Mariner* but of a different class. The most active element is *Mariner-1 D. elegans*, with 16 transfers (fig. 5). This TE has no transfers recorded from *D. elegans*, but is active in the *ananassae, ficusphila, chymomyza, quinaria, fenestrarum* (*Lordiphosa*), *leucophenga, miki* (*Lordiphosa*), *montium, obscura, testacea*, and *rhopaloa* subgroups. To our knowledge, the HT events in this transposon have been previously documented. We propose to name this TE *Minona*. The most dissimilar *Minona* elements present in different subgroups are still at least *>* 96% similar, thus the number of transfers would not be affected by changes to the 80-80-80 rule (fig. 5).

**Figure 5.**
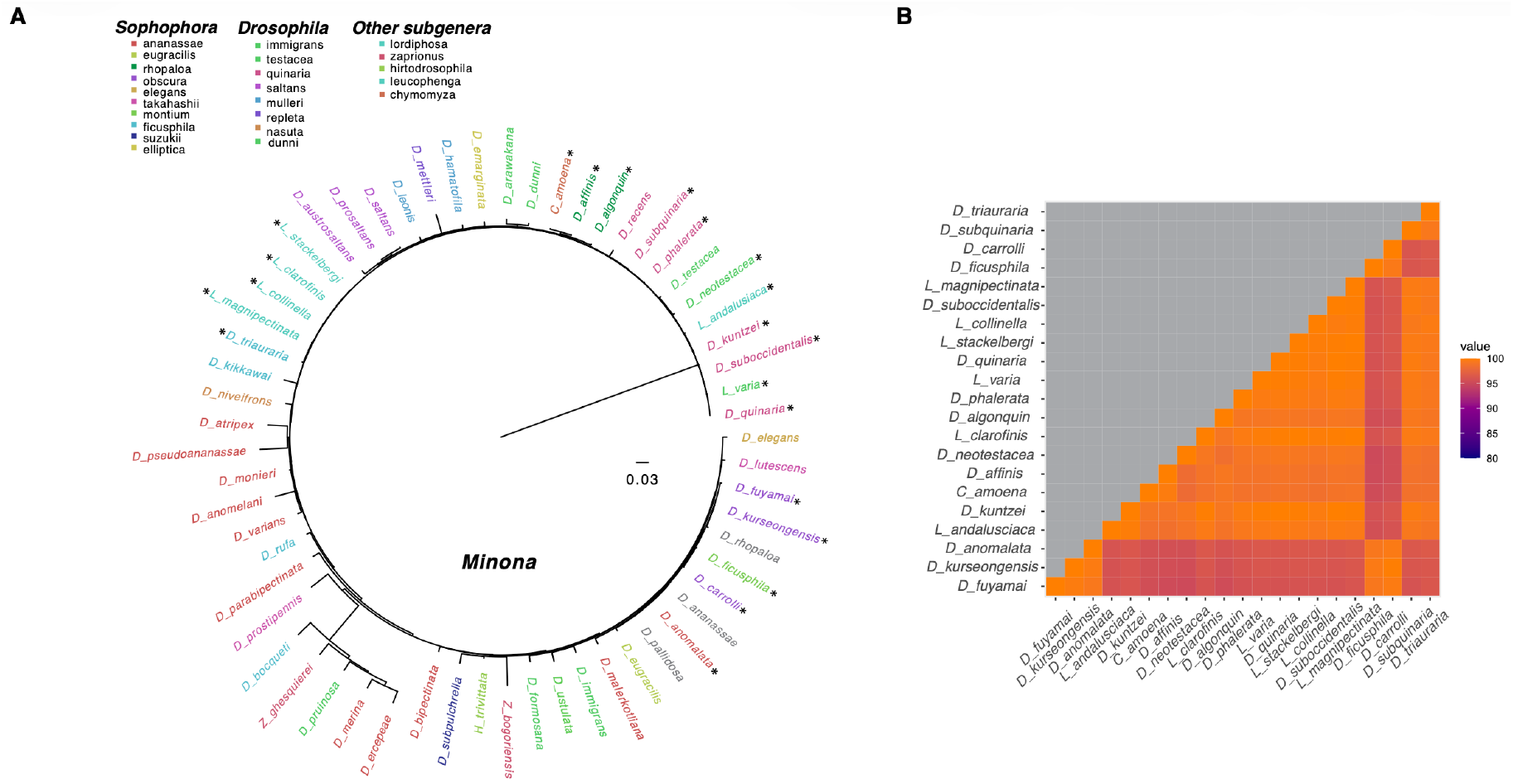
Phylogenetic tree of *Minona*. Each species name is colored according to its subgroup. *Zaprionus* species were not separated by subgroup in this figure but rather by genus. Gray species indicate that a transfer was documented in a relative too close to consider it an independent transfer event. For example, *D. ananassae, D. anomalata*, and *D. pallidosa* are all too similar to rule out inheritance by common descent, but a transfer into *D. pallidosa* is included. A star following the species name indicates a horizontal transfer events. B. A heatmap of similarity between TE sequences from species with a horizontal transfer event. Across all of the species with an active *Minona* element, the TE sequences are *>* 96% similar. In addition, similarity is not predicted by phylogenetic relationship - TE sequences from *Lordiphosa* species are more closely related to *D. kuntzei* and *D. triauraria* than *D. fuyamai* is to either.

Another TE with extensive HT is *Mariner-1 D. yakuba*, which is not active within most members of the *melanogaster* group, but is present in *D. yakuba* and *D. teissieri* (fig. 6). It has also transferred between 10 species of the *Zaprionus* group, *Leucophenga varia, Scaptodrosophila latifasciaeformis*, and two members of the *ananassae* subgroup *D. merina* and *D. ercepeae*. Increasing the threshold to 98.5% does not change the results. We propose to name this TE *Hatov* (fig. 6). The transfer between *D. merina* and *D. erecepeae* would be attributed to a different TE if we were to raise the required similarity between all TEs to 90%. We have also named three additional TEs in this dataset - *Sojourner, Judii*, and *Fizzle* - which corresponded to more than one other generically named TE and were renamed to reduce confusion (i.e. *Fizzle* corresponds to three other TEs in Repbase with different names consisting of *gypsy* and a number).

**Figure 6.**
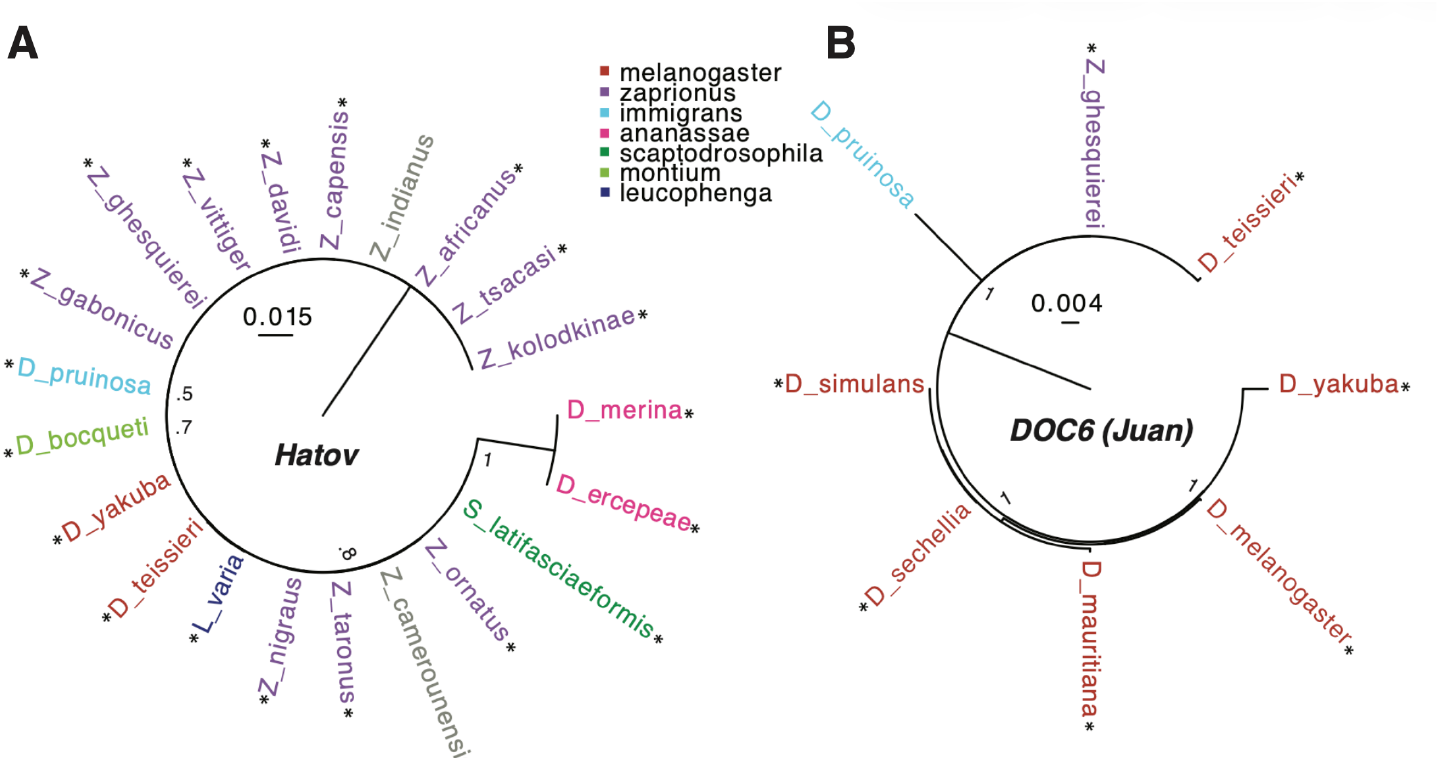
A-B. Phylogenetic trees showing the relationship between TEs in different species of dipterans. Note that *Zaprionus* is collapsed into a single group, although *Z. ghesqueirei* is a member of the *inermis* group while the remainder are *vittiger* group. Of the *Drosophila, D. pruinosa* is a member of the *Drosophila* subgenus while all others are *Sophophora*. The key indicates subgroup and is valid for both trees. A. *Hatov*, a TE that has been spreading primarily in African dipterans. HT events are indicated with an asterix. B. The second TE is *DOC6* (*Juan*) which has been documented spreading in the *melanogaster* group.

### Phylogenetic distance of horizontal transfer

We summarized the number of transfers between subgenera and subgroups.For example, we wanted to determine if LTRs move between different subgenera more commonly than non-LTRs. Due to the fact that many of these groups are potentially paraphyletic, the actual distance of the transfers is variable. However, in general it represents a group of TEs that move further than some other TEs. We found that DNA transposons move larger distances by far compared to LTRs and Non-LTRs (Figure 6). While LTRs make up the vast majority of transfers, DNA transpons are involved in 83% of between subgenera transfers and 62% of transfers between subgroups. Note that this result is not affected by the 80-80-80 rule - distance of transfer was summarized using only TEs that were *>* 99% similar. This could represent a separate evolutionary strategy, in which DNA transposons move less frequently to more naive genomes, potentially facilitating their establishment (Figure 6). This pattern is largely driven by *Mariner* elements, which make up 67% of transfers due to DNA transposons. This number is also likely an underestimate, as we assigned TEs transfer partners based on phylogenetic proximity.

The TE with some of the largest transfer distances is *POGO* (Figure 7). The *POGO* sequences present in different species are more than 98% similar across all genomes, thus this is not affected by the 80-80-80 rule. Within the *melanogaster* species group *POGO* has never been detected outside the *D. melanogaster*, a finding which is recapitulated here [Tudor et al., 1992]. Its presence in multiple species of *Zaprionus* suggests a likely origin in that genus, but more species of *Scaptodrosophila* and *Leucophenga* would need to be sequenced to say conclusively.

**Figure 7.**
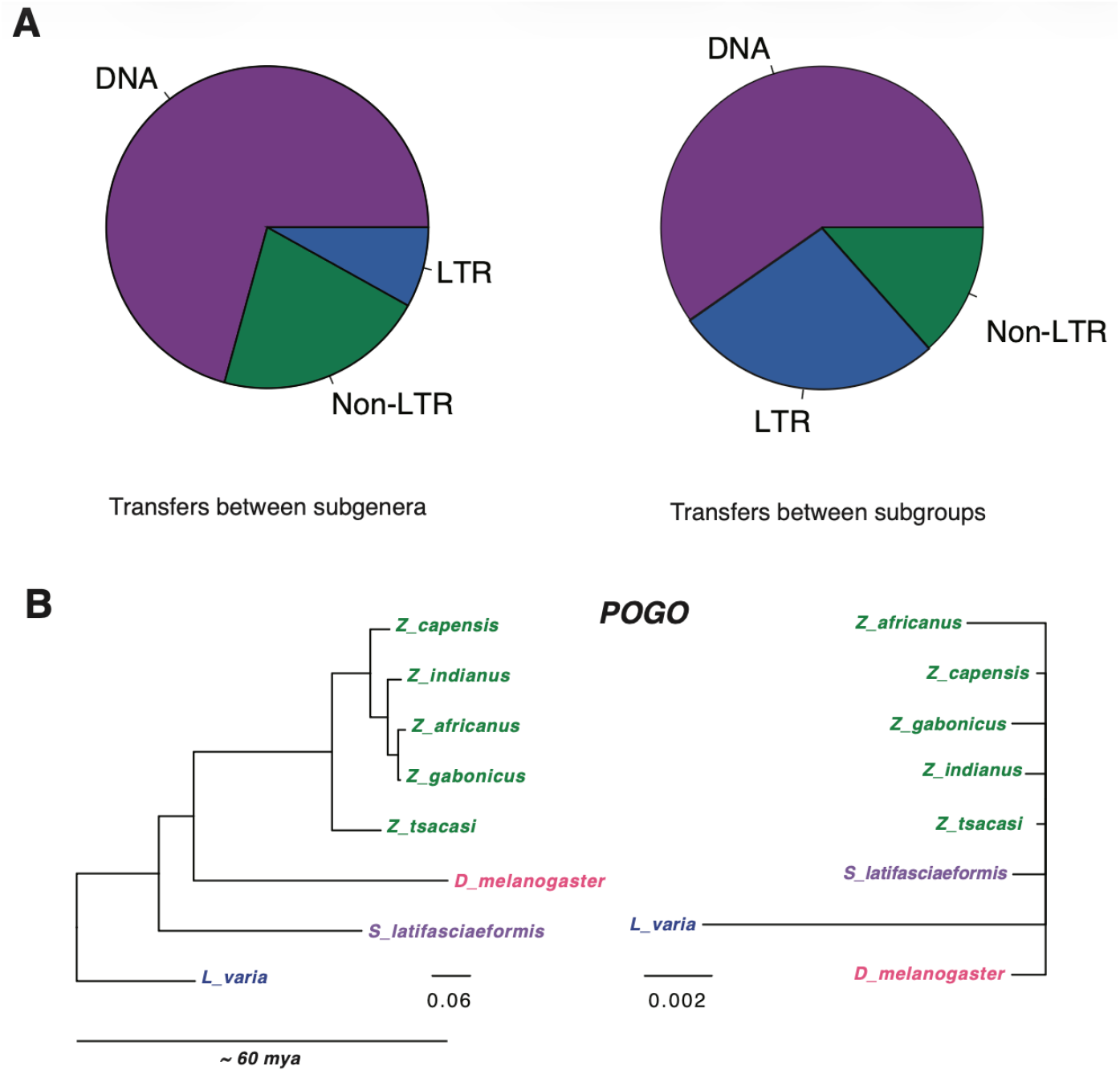
**A.** The proportion of TEs transferring between subgenera that belong to different TE types. **B**. The proportion of TEs transferring between subgroups that belong to different TE types. **C**. Phylogenetic tree of the species whose genomes harbor *POGO* using distances from Drosophyla, and a tree of *POGO* itself (right hand side). Note the large difference in scale between the two trees. The *POGO* phylogeny is essentially a star, with no differentiation between the majority of the species.

### HT in *D. melanogaster*

50 of the recent TE transfers involve *D. melanogaster* and have largely been documented elsewhere, for example *opus, Burdock*, and *Max*-element [Vieira et al., 1999, Pianezza and Kofler, 2025, Scarpa et al., 2023, Signor et al., 2023]. Less well documented transfers include *Chimpo, BEL* (also known as *3S18*), and *Blastopia* (also known as *flea*) [Bargues and Lerat, 2017]. *D. melanogaster* also either donated or acquired the *POGO* element from *Z. capensis* and *S*.*latifasciaformis*, which was not previously documented (Figure 7).

Aside from *POGO* (and the well documented *P*-element), species outside the *melanogaster* complex were not frequent donors and/or receivers with *D. melanogaster*, however *D. merina* likely donated or received *Transpac* and *ACCORD*, while *D. vallismaia* was likely involved in HT of *Bari*. In addition, *Z. tscacasi* likely donated or received *DOC2* from *D. melanogaster*. *D. vallismaia* and *D. merina* are both members of the *ananassae* group. *D. merina* is native to Madagascar, while *D. vallismaia* is native to the Seychelles. The relative importance of *D. merina* and other members of that subgroup such as *D. ercepeae* in HT with the *melanogaster group* is surprising, and has been noted elsewhere [Pianezza and Kofler, 2025].

### Abundant HT in other species

At the level of subgroup, as expected, *melanogaster* has the most transfers at 159. The *D. melanogaster* genome has been extensively analyzed for TE content, and all of those TEs are in this dataset, so the dataset is inherently biased towards detecting transfers in the *melanogaster* group. However, the *rhopaloa* subgroup has 56 annotated HT events. There are also 46 within the *bipectinata* species complex, a subset of the *ananassae* subgroup. While the abundance of HT in the *melanogaster* subgroup is undoubtedly due to the level of scrutiny the group has received, the *bipectinata* and *rhopaloa* groups have not been extensively annotated. This demonstrates that HT is occurring at high levels outside of *D. melanogaster*, even in species that are not considered human commensals. The *bipectinata* complex is of particular interest here, as we did not allow transfers between members of the *bipectinata* complex: *D. bipectinata, D. malerkotliana*, or *D. parabipectinata* because they are suspected (but not known) to still be interbreeding [Kopp and Barmina, 2005]. We did allow transfers from these species to *D. pseudoananassae* or any other species. Even restricting transfers to outside the complex, we still recorded 46 HT events involving this complex. An example of a TE with transfers in the *bipectinata* complex is shown in fig. 8, which includes transfers between species 36 mya diverged. The vast majority of HT events including the bipectinata complex are from the three species in the complex to *D. pseudoananassae* (34), which do not inter-breed. Other species involved in HT included *D. atripex, D. anomelani, D. eugracilis, D. birchii, D. biarmipes, D. ananassae, D. rufa*, and *D. varians*. It is unclear why the *bipectinata* complex would be such a hotspot for horizontal transfer - its TEs are no more well annotated than any other non-*melanogaster* group in the dataset.

**Figure 8.**
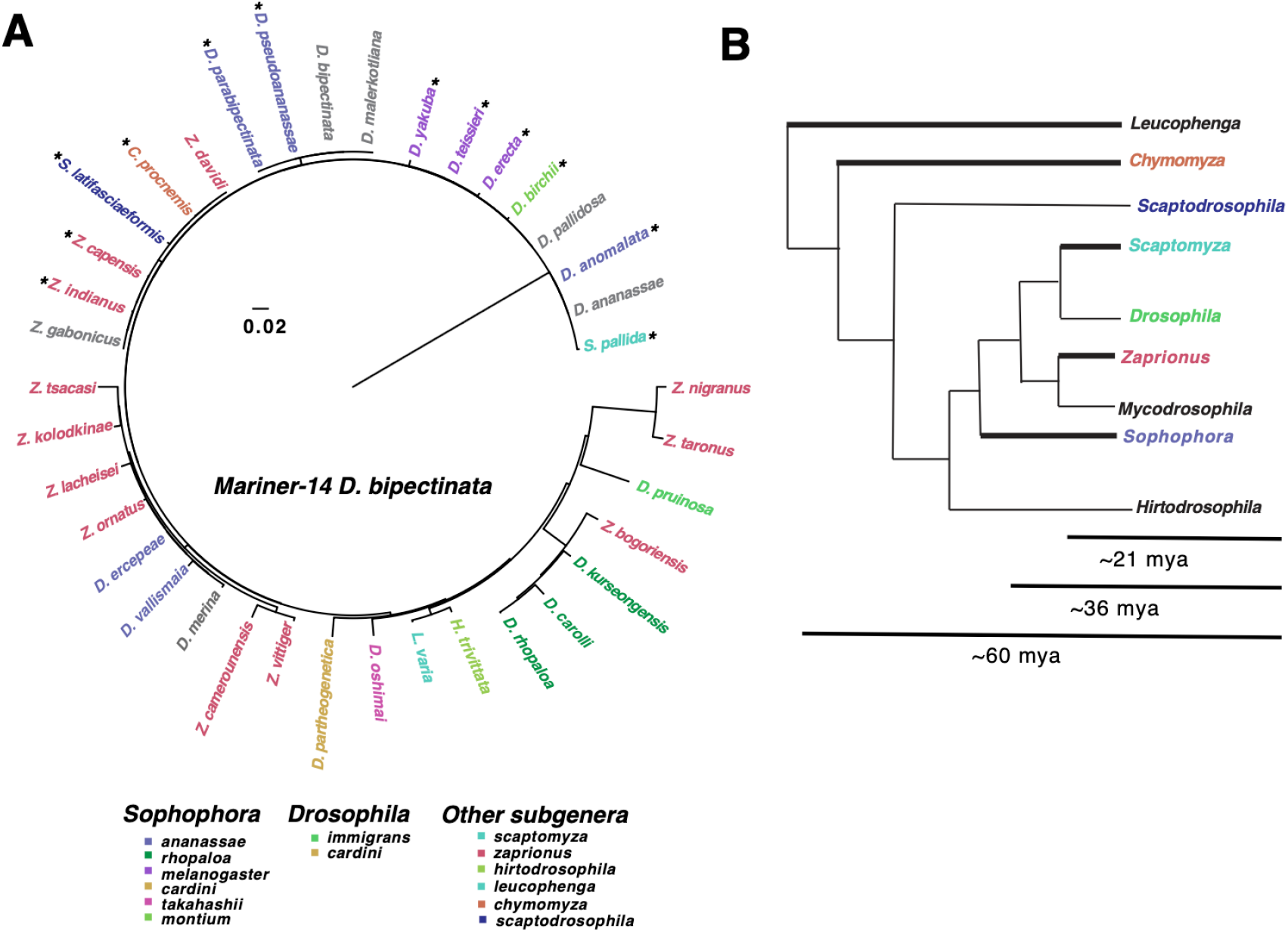
**A** HT in *Mariner-14 D. bipectinata*. Subgroup is indicated by color. This is essentially a star phylogeny thus the ordering of species is not necessarily reflective of the interrelationship between TEs. They are too similar in most cases for the exact relationships to be discerned. Species which were too closely related for HT to be included are shown in gray. B. A tree of dipterans included in the *Mariner-14 D. bipectinata* phylogeny. Lineages where a recent HT has occurred are indicated by thicker lines on the phylogeny. *Sophophora* is colored according to the most frequent group, *ananassae. Drosophila* is colored for *immigrans. Mariner-14 D. bipectinata* has transferred recently in *Chymomyza, Scaptodrosophila, Zaprionus, Scaptomyza, Drosophila*, and *Sophophora*. This represents a spread across more than 36 mya of indepedent evolution

## Discussion

In this paper we set out to track recent invasions of TEs across almost 400 *Drosophila* genomes (as well as close relatives). We detected 648 recent invasions of 268 different TEs, filtering for TE sequences that were *>* 99% similar. LTRs were responsible for the majority of HT events, however these invasions typically occurred over shorter evolutionary distances than invasions by DNA elements [Peccoud et al., 2017]. This suggests that the two types of TEs may have different invasion strategies. For example, LTRs may move more quickly but be restricted to more similar genomes, while DNA transposons are more flexible with regard to the genome but move larger phylogenetic distances less frequently. Some DNA transposons do only require their transposase to transpose in vitro [Lampe et al., 1996, Hartl et al., 1997]. Previously, insects were sampled at much higher phylogenetic distances and it was proposed that DNA elements engage in HT the most frequently - here we have evidence that it is dependent upon the phylogenetic distance of the species investigated [Peccoud et al., 2017, Silva et al., 2004]. Larger phylogenetic distances may ensure that the genome is naive with regard to the TE, compared to a close relative which may have already encountered a similar TE. Invasion of a similar TE would trigger the piRNA pathway to suppress the TE as it attempted to invade [Gainetdinov et al., 2023]. It is also possible that they use different vectors to spread, some of which are less specialized to a particular genus of drosophilid [**???**]. *Mariner* elements are known to be phylogenetically widespread and active in many different species [Capy et al., 1990, Lohe et al., 1995, Robertson, 1993, Wallau et al., 2014].

An interesting future direction would be to compare copy number between the different types of TEs, as this could also reflect different evolutionary strategies between TE types.

Many factors are going to bias the distribution of HT that we uncover - for example, the number of species within a group that is sequenced or the number of annotated TEs that originated from that group. For example, given the propensity for TEs to spread between close relatives it is more likely that you will detect an HT event if more species within a group have been sequenced. The relationship between TEs annotated in a group and the likelihood of finding HT is less clear. It was very common for a TE named after a species to be active primarily in a completely different group. For example, while *Mariner-1 D. elegans* (now *Minona*) was present in *D. elegans*, it did not engage in HT recently from this species. It did, however, engage in 16 transfers in other species. This is an extreme example, but it goes to show that while bias in TE annotation (for example, the focus on *D. melanogaster*) will impact the distribution of HT we find, it is not as clear cut a relationship as one might expect.

Our summary of the number of HT events between subgroups does represent very different phylogenetic distances - for example a jump between the *ananassae* subgroup and *melnaogaster* subgroup is much different than a jump between *Leucophenga* and *melanogaster*. Regardless, it is a useful categorization of transfers between quite closely related species and those further away. The way in which we assigned transfer partners was also based on assumptions that are not always true - we prioritized within subgroup transfers over between subgroup transfers and both over between-subgenera transfers. It is entirely possible that the rate of transfer between different subgenera is higher, and has happened multiple times for a single TE (i.e. multiple transfers between *Drosophila* and *Zaprionus* rather than a single transfer followed by within-*Zaprionus* transfers). For the vast majority of the HT events categorized in this paper, there is no way to reliably distinguish transfer partners given the high sequence similarity between TEs.

We also find a high rate of recent HT in quite a number of subgroups in addition to *D. melanogaster*. This suggests that high rates of HT may not be unique to *D. melanogaster* [Scarpa et al., 2023, Pianezza et al., 2023, 2025]. It has been suggested that HT has been accelerated by human movement, which is likely, but perhaps human movement of different drosophilids is extensive enough that it is not limited by the known distribution of the species i.e. the majority of species are essentially cosmopolitan. It is also possible that it is movement of the species pathogens that is impacting HT (if that is a vector for HT), as new pathogens might spread more readily than new flies. We do not yet know if this rate of invasion is historical for species not considered human commensals, but this is an interesting open question.

A thorough understanding of HT between species has been hindered by the lack of complete assemblies of repetitive regions, which was not possible until the advent of long-read sequencing [van Dijk et al., 2023]. The pace at which this data is now being produced is creating a unique opportunity to understand the dynamics of repetitive elements. However, it can still be difficult to place one’s results in the context of existing literature. For example, one issue that has hindered our understanding of HT is confusion over the level at which HT is detected in many older manuscripts - for example if *Mariner* is detected across insect species, does this refer to the class, a group within *Mariner*, or an individual element. Another is naming, TEs are frequently given generic names such as *gypsy-5* through *50*, followed by a species name, and it can be difficult to differentiate different elements (and there are many repeats). TE sequences are also frequently not deposited in any central resource such as NCBI or Dfam. In some cases, the sequences referred to in a manuscript cannot be uncovered at all. Improved central databases and naming conventions will help ameliorate these problems as more TEs are discovered. Approaches which emphasize reference-free TE discovery will also be of considerable help in this regard.

Many of the TEs in the original dataset were not found to have undergone recent HT. However, our cutoff of *>* 99% similarity was somewhat arbitrary and does not necessarily imply that they have not been active in the somewhat recent past. A subsequent study focusing on estimating genome divergence relative to TE divergence will be a welcome contribution to our understanding of HT in the recent past. In addition, investment in museum sequencing will contribute to our understanding of the rate of TE invasion in these other species [Scarpa et al., 2023]. There are still many unanswered questions about HT in dipterans - for example how does a new TE invasion impact the population of flies? In one study, we found that the population genetics of *D. sechellia* was very different on islands that had been invaded by two TEs versus islands that had not [Scarpa et al., 2025]. Islands that had been invaded by the two TEs had flies that were all very closely related to each other, while islands without the two TEs had more population structure [Scarpa et al., 2025]. Can TE invasions cause population bottlenecks? Is there selection for TE insertions within TE silencing regions, or are insertions into these regions abundant as has been suggested for the *P* –element [Wang et al., 2023]? How do the invasion dynamics of TEs with envelope proteins differ from those that do not have envelope proteins [Senti et al., 2025]? While many questions remain about the dynamics and genomic consequences of TE invasions, it is clear that they are a powerful force shaping the evolution of species genomes.

## Acknowledgments

SS would like to thank F. Belt, S. & F. Emery, and P. Senn for inspiration during the production of this manuscript. SP would like to thank his late grandma Shanti Devi for helping him learn how to become a better human being.

## Author contributions

S.S. conceived the project. S.P. performed computational analyses; S.S. performed manual curation. S.P. and S.S. co-wrote the manuscript.

## Funding

This work was supported by the National Science Foundation Established Program to Stimulate Competitive Research grants NSF-EPSCoR-1826834 and NSF-EPSCoR-2032756 to SS. In addition this work was funding by the National Institutes of Health R35GM155272 to SS.

## Conflicts of Interest

The author(s) declare(s) that there is no conflict of interest regarding the publication of this article.

## Data Availability

The genomes used in this study are all publicly available from our cited sources, as are the TE sequences used [**???**]. Newly named TEs such as *Minona* were deposited in Dfam. The processed data discussed in this manuscript is all included in the supplemental files.

## Notes

### Competing Interest Statement

The authors have declared no competing interest.

### Summary of Updates

The main purpose of the revision was to alter the analysis to focus more on subgenus and subgroup rather than individual transfer events, and to incorporate additional phylogenetic information about subgenera.

## References

A. A. Aravin, G. J. Hannon, and J. Brennecke. The Piwi-piRNA pathway provides an adaptive defense in the transposon arms race. Science, 318(5851):761–764, 2007.

N. Bargues and E. Lerat. Evolutionary history of ltr-retrotransposons among 20 drosophila species. Mobile DNA, 8:1–15, 2017.

J. Brennecke, A. A. Aravin, A. Stark, M. Dus, M. Kellis, R. Sachidanandam, and G. J. Hannon. Discrete small RNA-generating loci as master regulators of transposon activity in *Drosophila*. Cell, 128(6):1089–1103, 2007.

J. Brennecke, C. D. Malone, A. A. Aravin, R. Sachidanandam, A. Stark, and G. J. Hannon. An epigenetic role for maternally inherited piRNAs in transposon silencing. Science, 322(5906):1387–1392, 2008.

C. Camacho, G. Coulouris, V. Avagyan, N. Ma, J. Papadopoulos, K. Bealer, and T. L. Madden. Blast+: architecture and applications. BMC bioinformatics, 10(1):421, 2009.

P. Capy, F. Chakrani, F. Lemeunier, D. Hartl, and J. David. Active mariner transposable elements are widespread in natural populations of drosophila simulans. Proceedings of the Royal Society of London. Series B: Biological Sciences, 242(1303):57–60, 1990.

M. Chakraborty, C. Chang, D. Khost, J. A. J Vedanayagam, Y. Liao, K. Montooth, C. Meiklejohn, A. Larracuente, and J. Emerson. Evolution of genome structure in the *Drosophila simulans* species complex. Genome Research, 31:380–396, 2021.

S. B. Daniels, K. R. Peterson, L. D. Strausbaugh, M. G. Kidwell, and A. Chovnick. Evidence for horizontal transmission of the P transposable element between *Drosophila* species. Genetics, 124(2):339–55, 1990.

N. de la Chaux and A. Wagner. Evolutionary dynamics of the ltr retrotransposons roo and rooa inferred from twelve complete drosophila genomes. BMC evolutionary biology, 9(1):205, 2009.

G. R. Dias, E. G. Dupim, T. Vanderlinde, B. Mello, and A. B. Carvalho. Timing and pattern of early diversification in drosophilidae (diptera). Molecular Biology and Evolution, 42(11):msaf269, 2025.

D. G. Eickbush and T. H. Eickbush. Vertical transmission of the retrotransposable elements R1 and R2 during the evolution of the *Drosophila melanogaster* species subgroup. Genetics, 139(2):671–684, 1995.

C. Feschotte and E. J. Pritham. DNA transposons and the evolution of eukaryotic genomes. Annual review of genetics, 41:331–68, 2007.

M. Fraser, J. S. Brusca, G. E. Smith, and M. D. Summers. Transposon-mediated mutagenesis of a baculovirus. Virology, 145(2):356–361, 1985.

I. Gainetdinov, J. Vega-Badillo, K. Cecchini, A. Bagci, C. Colpan, D. De, S. Bailey, A. Arif, P.-H. Wu, I. J. MacRae, et al. Relaxed targeting rules help piwi proteins silence transposons. Nature, 619(7969):394–402, 2023.

L. S. Gunawardane, K. Saito, K. M. Nishida, K. Miyoshi, Y. Kawamura, T. Nagami, H. Siomi, and M. C. Siomi. A slicer-mediated mechanism for repeat-associated siRNA 5’ end formation in *Drosophila*. Science, 315(5818):1587–1590, 2007.

D. L. Hartl, A. R. Lohe, and E. R. Lozovskaya. Modern thoughts on an ancyent marinere: function, evolution, regulation. Annual review of genetics, 31(1):337–358, 1997.

B. D. Heath, R. D. Butcher, W. G. Whitfield, and S. F. Hubbard. Horizontal transfer of wolbachia between phylogenetically distant insect species by a naturally occurring mechanism. Current Biology, 9(6):313–316, 1999.

M. A. Houck, J. B. Clark, K. R. Peterson, and M. G. Kidwell. Possible horizontal transfer of drosophila genes by the mite proctolaelaps regalis. Science, 253(5024):1125–1128, 1991.

J. P. Huelsenbeck and F. Ronquist. Mrbayes: Bayesian inference of phylogenetic trees. Bioinformatics, 17 (8):754–755, 2001.

J. Jehle, E. Fritsch, A. Nickel, J. Huber, and H. Backhaus. Tcl4. 7: a novel lepidopteran transposon found in cydia pomonella granulosis virus. Virology, 207(2):369–379, 1995.

K. Katoh and H. Toh. Recent developments in the mafft multiple sequence alignment program. Briefings in bioinformatics, 9(4):286–298, 2008.

K. Katoh, K. Misawa, K.-i. Kuma, and T. Miyata. Mafft: a novel method for rapid multiple sequence alignment based on fast fourier transform. Nucleic acids research, 30(14):3059–3066, 2002.

K. Katoh, G. Asimenos, and H. Toh. Multiple alignment of dna sequences with mafft. In Bioinformatics for DNA sequence analysis, pages 39–64. Springer, 2009.

B. Y. Kim, J. R. Wang, D. E. Miller, O. Barmina, E. Delaney, A. Thompson, A. A. Comeault, D. Peede, E. R. D’Agostino, J. Pelaez, J. M. Aguilar, D. Haji, T. Matsunaga, E. E. Armstrong, M. Zych, Y. Ogawa, M. Stamenković-Radak, M. Jelić, M. S. Veselinović, M. Tanasković, P. Erić, J.-J. Gao, T. K. Katoh, M. J. Toda, H. Watabe, M. Watada, J. S. Davis, L. C. Moyle, G. Manoli, E. Bertolini, V. Košťál, R. S. Hawley Takahashi, C. D. Jones, D. K. Price, N. Whiteman, A. Kopp, D. R. Matute, and D. A. Petrov. Highly contiguous assemblies of 101 drosophilid genomes. eLife, 10:e66405, 2021.

Y. Kim, H. R. Gellert, S. H. Church, A. Suvorov, S. S. Anderson, O. Barmina, S. G. Beskid, A. A. Comeault, K. Nicole Crown, S. E. Diamond, S. Dorus, T. Fujichika, J. A. Hemker, J. Hrcek, M. Kankare, T. Katoh, K. N. Magnacca, R. A. Martin, T. Matsunaga, M. J. Medeiros, D. E. Miller, S. Pitnick, S. Simoni, T. E. Steenwinkel, M. Schiffer, Z. A. Syed, A. Takahashi, K. H.-C. Wei, T. Yokoyama, M. B. Eisen, A. Kopp, D. Matute, D. J. Obbard, P. M. O’Grady, D. K. Price, M. J. Toda, T. Werner, and D. A. Petrov. Single-fly assemblies fill major phylogenomic gaps across the drosophilidae tree of life. bioRxiv, Oct. 2023. doi: 10.1101/2023.10.02.560517. URL http://dx.doi.org/10.1101/2023.10.02.560517.

R. Kofler, T. Hill, V. Nolte, A. Betancourt, and C. Schlötterer. The recent invasion of natural *Drosophila simulans* populations by the P-element. PNAS, 112(21):6659–6663, 2015a.

R. Kofler, V. Nolte, and C. Schlötterer. Tempo and mode of transposable element activity in *Drosophila*. PLoS Genetics, 11(7):e1005406, 2015b.

N. Kondo, N. Nikoh, N. Ijichi, M. Shimada, and T. Fukatsu. Genome fragment of wolbachia endosymbiont transferred to x chromosome of host insect. Proceedings of the National Academy of Sciences, 99(22): 14280–14285, 2002.

A. Kopp and O. Barmina. Evolutionary history of the drosophila bipectinata species complex. Genetics Research, 85(1):23–46, 2005.

D. J. Lampe, M. Churchill, and H. M. Robertson. A purified mariner transposase is sufficient to mediate transposition in vitro. The EMBO journal, 15(19):5470–5479, 1996.

A. R. Lohe, E. N. Moriyama, D.-A. Lidholm, and D. L. Hartl. Horizontal transmission, vertical inactivation, and stochastic loss of mariner-like transposable elements. Molecular biology and evolution, 12(1):62–72, 1995.

C. D. Malone, J. Brennecke, M. Dus, A. Stark, W. R. McCombie, R. Sachidanandam, and G. J. Hannon. Specialized piRNA pathways act in germline and somatic tissues of the *Drosophila* ovary. Cell, 137(3): 522–535, 2009.

S. B. Marion, K. Focht, I. Hamid, E. S. Iversen, H. John, B. Manzano-Winkler, A. Navarra, S. Pangare, M. Zarei, and M. A. Noor. Transposable elements contribute substantially to naturally occurring genetic lethality in drosophila melanogaster. PLoS biology, 24(3):e3003467, 2026.

K. Maruyama and D. Hartl. Evolution of the transposable element mariner in drosophila species. Genetics, 128(2):319, 1991a.

K. Maruyama and D. L. Hartl. Evidence for interspecific transfer of the transposable element mariner between drosophila and zaprionus. Journal of molecular evolution, 33(6):514–524, 1991b.

V. Mérel, M. Boulesteix, M. Fablet, and C. Vieira. Transposable elements in drosophila. Mobile DNA, 11 (1):23, 2020.

D. W. Miller and L. K. Miller. A virus mutant with an insertion of a copia-like transposable element. Nature, 299(5883):562–564, 1982.

L. Modolo, F. Picard, and E. Lerat. A new genome-wide method to track horizontally transferred sequences: application to *Drosophila*. Genome biology and evolution, 6(2):416–432, 2014.

F. F. Nascimento, M. D. Reis, and Z. Yang. A biologist’s guide to Bayesian phylogenetic analysis. Nature Ecology & Evolution, 1(10):1446–1454, Sept. 2017. ISSN 2397-334X. doi: 10.1038/s41559-017-0280-x. URL https://www.nature.com/articles/s41559-017-0280-x.

E. Paradis and K. Schliep. ape 5.0: an environment for modern phylogenetics and evolutionary analyses in r. Bioinformatics, 35(3):526–528, 2019.

E. Paradis, J. Claude, and K. Strimmer. Ape: analyses of phylogenetics and evolution in r language. Bioinformatics, 20(2):289–290, 2004.

J. Peccoud, V. Loiseau Cordaux, and C. Gilbert. Massive horizontal transfer of transposable elements in insects. Proc Natl Acad Sci U S A, 114(18):4721–26, 2017.

R. Pianezza and R. Kofler. Biogeography shapes the te landscape of drosophila melanogaster. bioRxiv, 2025. doi: 10.1101/2025.05.22.655554. URL https://www.biorxiv.org/content/early/2025/05/27/2025.05.22.655554.

R. Pianezza, A. Scarpa, P. Narayanan, S. Signor, and R. Kofler. Spoink, a ltr retrotransposon, invaded D. melanogaster populations in the 1990s. bioRxiv, 2023.

R. Pianezza, A. Scarpa, A. Haider, S. Signor, and R. Kofler. Spatio-temporal tracking of three novel transposable element invasions in drosophila melanogaster over the last 30 years. Molecular Biology and Evolution, page msaf143, 2025.

P. Rice, I. Longden, and A. Bleasby. Emboss: the european molecular biology open software suite. Trends in genetics, 16(6):276–277, 2000.

H. M. Robertson. The mariner transposable element is widespread in insects. Nature, 362(6417):241–245, 1993.

F. Ronquist, M. Teslenko, P. Van Der Mark, D. L. Ayres, A. Darling, S. Höhna, B. Larget, L. Liu, M. A. Suchard, and J. P. Huelsenbeck. Mrbayes 3.2: efficient bayesian phylogenetic inference and model choice across a large model space. Systematic biology, 61(3):539–542, 2012.

A. Scarpa, R. Pianezza, F. Wierzbicki, and R. Kofler. Genomes of historical specimens reveal multiple invasions of ltr retrotransposons in *Drosophila melanogaster* populations during the 19th century. PNAS, 2023.

A. Scarpa, R. Pianezza, H. R. Gellert, A. Haider, B. Y. Kim, E. C. Lai, R. Kofler, and S. Signor. Double trouble: two retrotransposons triggered a cascade of invasions in drosophila species within the last 50 years. Nature Communications, 16(1):516, 2025.

A. S. Seetharam and G. W. Stuart. Whole genome phylogeny for 21 drosophila species using predicted 2b-rad fragments. PeerJ, 1:e226, 2013.

K.-A. Senti and J. Brennecke. The piRNA pathway: a fly’s perspective on the guardian of the genome. Trends in genetics, 26(12):499–509, 2010.

K.-A. Senti, B. Rafanel, D. Handler, C. Kosiol, C. Schlötterer, and J. Brennecke. Co-evolving infectivity and expression patterns drive the diversification of endogenous retroviruses. The EMBO Journal, pages 1–20, 2025.

F. Sievers and D. G. Higgins. Clustal omega for making accurate alignments of many protein sequences. Protein Science, 27(1):135–145, 2018.

S. Signor, J. Vedanayagam, B. Kim, F. Wierzbicki, R. Kofler, and E. Lai. Rapid evolutionary diversification of the flamenco locus across simulans clade *Drosophila* species. PLoS Genet, 19, 2023.

J. C. Silva, E. L. Loreto, and J. B. Clark. Factors that affect the horizontal transfer of transposable elements. Current issues in molecular biology, 6:57–71, 2004.

S. U. Song, T. Gerasimova, M. Kurkulos, J. D. Boeke, and V. G. Corces. An env-like protein encoded by a drosophila retroelement: evidence that gypsy is an infectious retrovirus. Genes & development, 8(17): 2046–2057, 1994.

S. Srivastav, C. Feschotte, and A. G. Clark. Rapid evolution of piRNA clusters in the *Drosophila melanogaster* ovary. bioRxiv, 2023.

F. K. Teixeira, M. Okuniewska, C. D. Malone, R.-X. Coux, D. C. Rio, and R. Lehmann. pirna-mediated regulation of transposon alternative splicing in the soma and germ line. Nature, 552(7684):268–272, 2017.

M. Tudor, M. Lobocka, M. Goodell, J. Pettitt, and K. O’Hare. The pogo transposable element family of drosophila melanogaster. Molecular and General Genetics MGG, 232(1):126–134, 1992.

E. L. van Dijk, D. Naquin, K. Gorrichon, Y. Jaszczyszyn, R. Ouazahrou, C. Thermes, and C. Hernandez. Genomics in the long-read sequencing era. Trends in Genetics, 39(9):649–671, 2023.

S. Venner, V. Miele, C. Terzian, C. Biémont, V. Daubin, C. Feschotte, and D. Pontier. Ecological networks to unravel the routes to horizontal transposon transfers. PLOS Biology, 15(2):e2001536, 2017. ISSN 1545-7885. doi: 10.1371/journal.pbio.2001536.

C. Vieira, D. Lepetit, S. Dumont, and C. Biémont. Wake up of transposable elements following drosophila simulans worldwide colonization. Molecular biology and evolution, 16:1251–5, 10 1999. doi: 10.1093/oxfordjournals.molbev.a026215.

G. L. Wallau, P. Capy, E. Loreto, and A. Hua-Van. Genomic landscape and evolutionary dynamics of mariner transposable elements within the drosophila genus. BMC genomics, 15(1):727, 2014.

L. Wang, S. Zhang, S. Hadjipanteli, L. Saiz, L. Nguyen, E. Silva, and E. Kelleher. P-element invasion fuels molecular adaptation in laboratory populations of *Drosophila melanogaster*. Evolution, 77(4):980–994, 2023.

T. Wicker, F. Sabot, A. Hua-Van, J. L. Bennetzen, P. Capy, B. Chalhoub, A. Flavell, P. Leroy, M. Morgante, O. Panaud, et al. A unified classification system for eukaryotic transposable elements. Nature Reviews Genetics, 8(12):973–982, 2007.

M. Yoshiyama, Z. Tu, Y. Kainoh, H. Honda, T. Shono, and K. Kimura. Possible horizontal transfer of a transposable element from host to parasitoid. Molecular biology and evolution, 18(10):1952–1958, 2001.

