## Supplemental File1 for "Recent horizontal transfer of transposable elements in *Drosophila*"

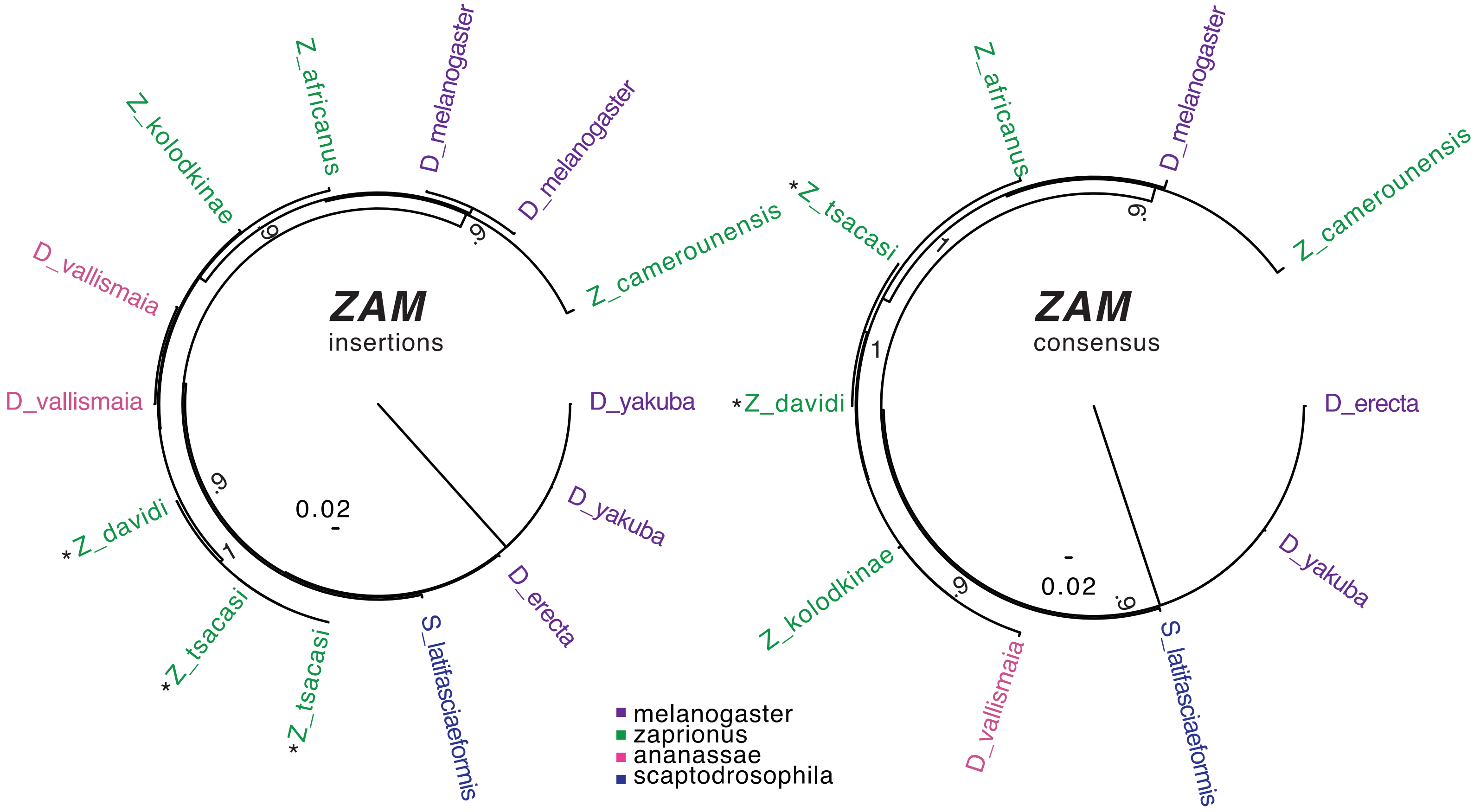


Supplemental File 1: An example of TE trees made with consensus sequences of *ZAM* and individual insertions. In some cases, i.e. *D. erecta*, only a single copy was available. While the ordering of the tree is not identical, overall the pattern of relationships is consistent. The portions of the tree with changes in ordering are essentially a star phylogeny. A second example is shown on the next page for *piggBac-2 Drosophila eugracilis*. The same overall pattern holds, in that the approach would not change the inference of horizontal transfer events.

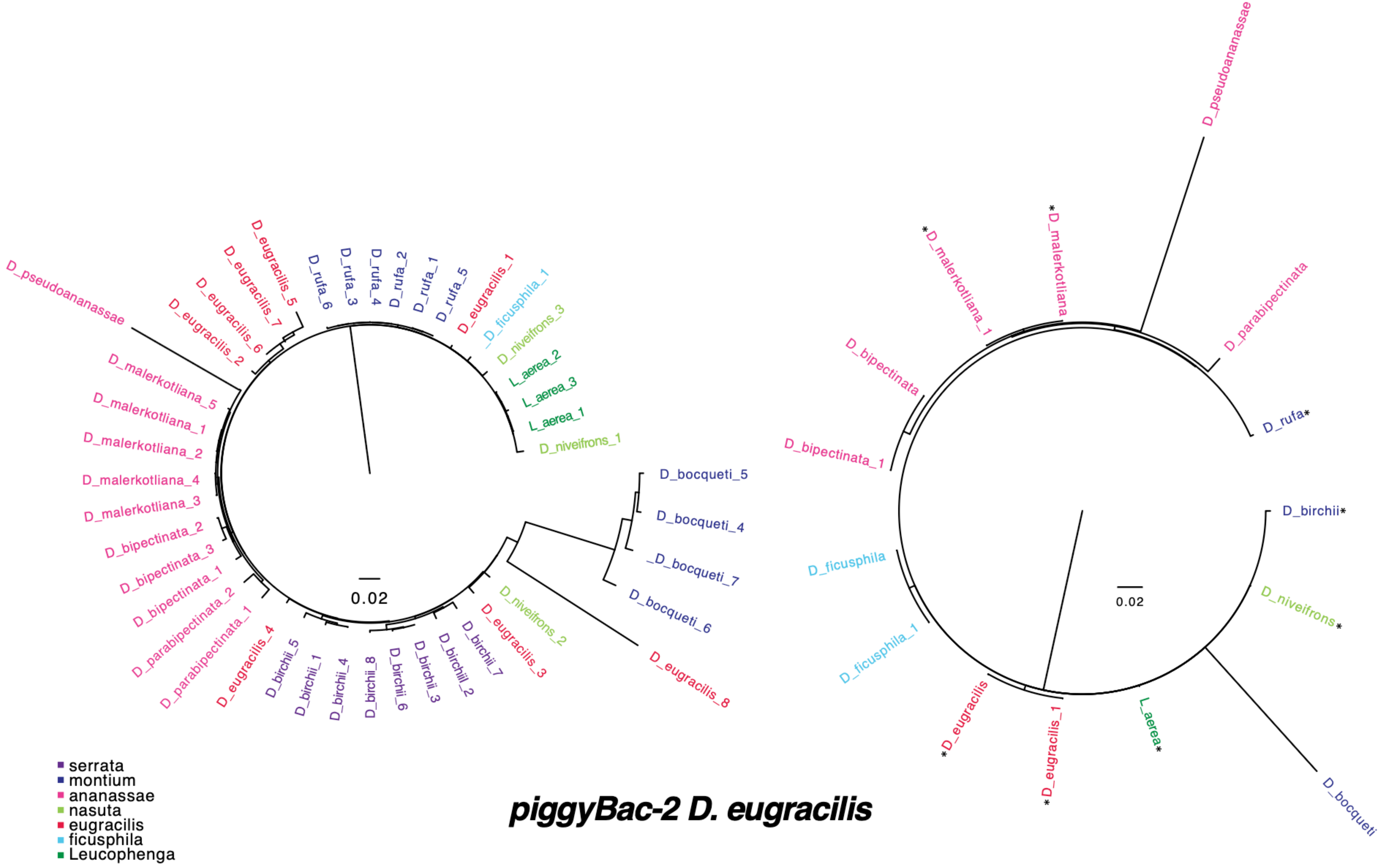


***piggyBac-2 D. eugracilis***
